# Socialization causes long-lasting behavioral changes

**DOI:** 10.1101/2024.04.25.591071

**Authors:** Beatriz Gil-Martí, Julia Isidro-Mézcua, Adriana Poza-Rodriguez, Gerson S. Asti Tello, Gaia Treves, Enrique Turiégano, Esteban J. Beckwith, Francisco A Martin

## Abstract

In modern human societies, social isolation acts as a negative factor for health and life quality. On the other hand, social interaction also has profound effects on animal and human behaviors, reducing aggressiveness, feeding and loss of sleep. Here, we observe that in the fly *Drosophila melanogaster* these behavioral changes long-last even after social interaction has ceased, suggesting that the socialization experience triggers behavioral plasticity. We find that impairing long-term memory mechanisms either genetically or by anesthesia abolishes the expected behavioral changes in response to social interaction. Furthermore, we show that socialization increases CREB-dependent neuronal activity and synaptic plasticity in the mushroom body, the main insect memory center analogous to mammalian hippocampus. We propose that social interaction triggers socialization awareness, understood as long-lasting changes in behavior caused by experience with mechanistic similarities to long-term memory formation.

## INTRODUCTION

Most animals live in social contexts. In our modern human society, the feeling of loneliness is increasing despite the technological advances in social media and communication^1^. The prolonged absence of social interaction has detrimental effects on quality of life, lifespan and several health problems^2,3^. In *Drosophila melanogaster*, social interaction strongly modulates several behaviors, diminishing male-to-male aggression, decreasing food consumption and, depending on the context, increasing or decreasing sleep, among others^4^. Socialization impacts several parallel modulatory systems^5^. In particular, activity-regulated genes in dopaminergic neurons modulate aggression and sleep in response to social enrichment^6–8^. Key clusters of dopaminergic neurons are also essential components of learning and memory circuits^9^, since they innervate the main *Drosophila* memory structure, the mushroom body (MB)^10^.

At the molecular level, long-term memory (LTM) formation in the MB requires *rutabaga (rut-adenylate cyclase)* and *d*unce (*dnc*-*cAMP phosphodiesterase)* gene functions, in order to adequately regulate cAMP levels and ensure neuronal plasticity^11^. cAMP signaling mediates CREB (c-AMP response binding element) phosphorylation. CREB is a conserved transcription factor that is key to form long-term memory and synaptic plasticity, among many other processes^11,12^. Social interaction causes structural changes in the MB, an effect that is abolished in mutant flies for memory-related genes like *rut* and *dnc*^13,14^. Furthermore, the function of such genes is necessary for immediate sleep changes triggered by social interaction^15,16^.

In this work, we inquired if socialization was able to generate long-lasting changes on behavior, and addressed how these changes are associated with synaptic plasticity. We showed that underlying mechanisms have similarities with LTM: an altered behavior in response to experience lasted for 8 or more hours after the training; it depended on cAMP levels and was blocked by anesthesia; and ultimately, it correlated with changes in number of CREB-responsive neurons and synapses. In summary, we propose that socialization awareness modifies long-term behavior sharing some underlying mechanisms that are characteristic of long-term memory processes.

## RESULTS

### LONG-TERM SOCIALIZATION-INDUCED FEEDING BEHAVIOR REQUIRES cAMP SIGNALING

Flies that experienced social interaction show reduced food consumption when compared with flies that were socially reared and posteriorly isolated^17^. We used single-fly CApillary FEeding -sCAFE-assay (modified from^18^) to extend these findings. We compared grouped flies with animals singly reared since eclosion, meaning that they were socially naive. As expected, there was a significant decrease in food uptake of 5-day socialized flies when compared to individual flies in the immediate 24 hours (0 h-24 h time window) (fig 1A). Next, to determine if such feeding effect is maintained even in the absence of social interaction, we slightly modified the socially-enriched paradigm: flies were group- or single-reared for 5 days and then animals from both experimental groups were kept isolated for additional 24 hours previous to assessing feeding (fig 1B). Using this protocol, we also detected a decreased food consumption of grouped flies in the 24 h-48 h time window, confirming a long-lasting effect of social interaction on feeding behavior (fig 1C). We reasoned that the most plausible candidate genes to play a role for such long-lasting effect would be memory-related genes, such as *rutabaga* (*rut*)^19^. Despite their past experience, isolated *rut* mutant flies in the 24 h-48 h period after socialization showed no differences in food intake with solitaire animals since eclosion (fig 1C). Besides, *rut* mutant flies do not change their feeding behavior during the first 24 hours (0 h-24 h), suggesting a requirement of cAMP for this response (fig 1A). To confirm the involvement of cAMP signaling we repeated the sCAFE assay in animals mutant for *dunce* (*dnc*). Results were comparable to *rut* mutant: *dnc* mutant flies fail to modify their food consumption not only during the first 24 hours after socialization (0 h-24 h) but also according to previous experience, in the 24 h-48 h period (fig 1A, C). Strikingly, feeding behavior of memory-related mutant animals lies in an intermediate state between socialized and isolated flies, maybe suggesting that their basal food consumption is different from wild type strain (see below). We propose that this long-lasting effect of social interaction on (feeding) behavior indicates a that animals have a socialization awareness that lasts even after social interaction has ceased.

**Figure 1.**
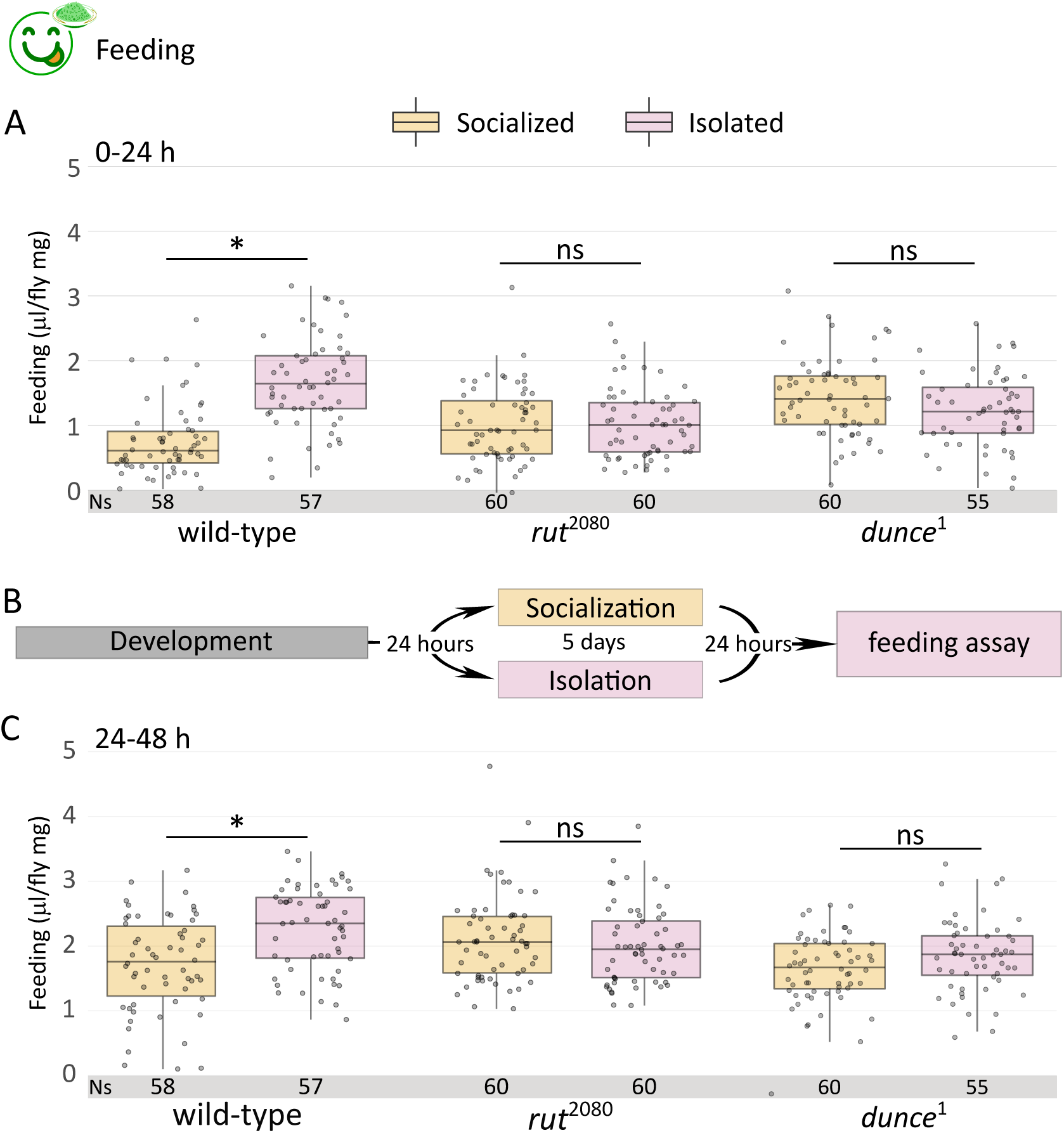
Reduced food consumption induced by socialization depends on memory-related genes. (A) Quantification of food consumption of *wt*, *rut* and *dnc* mutant flies in socialized and isolated conditions (single fly CAFE assay) in the 0-24 h time window (Kruskal-Wallis chi-squared = 75.905, df = 5, p-value = 6.022e-15; *post hoc* Dunn comparisons: wt^social^|wt^isolated^ p = 6.24e-13, *rut*^social^|*rut*^isolated^ p = 1.00, *dnc*^social^|*dnc*^isolated^ = 1.00). (B) Scheme of the socialization protocol: recently eclosed animals were either grouped or isolated for five days, and subsequently isolated for additional 24 hours before tested. C) Quantification of food consumption of *wt*, *rut* and *dnc* mutant flies in socialized and isolated conditions (sCAFE) in the 24 h-48 h h time window (Kruskal-Wallis chi-squared = 32.698, df = 5, p-value = 4.32e-06; *post hoc* Dunn comparisons: wt^social^|wt^isolated^ p = 1.04e-03, *rut*^social^|*rut*^isolated^ p = 1.00, and, in fig S1, *dnc*^social^|*dnc*^isolated^ = 1.00).

### LONG-TERM EFFECT OF SOCIALIZATION ON SLEEP IN ABSENCE OF INTERACTION

Sleep (particularly daytime sleep) is also regulated after social interaction. When flies are kept in mixed-sex groups immediately at eclosion, sleep increases^15^. However, courtship experience inhibits sleep in male flies ^20,21^, which lasts for several hours, proving a complex regulation of sleep by social cues and experiences. Most published sleep studies employ *Drosophila* Activity Monitors (DAMs), which only detect movement when the fly crosses a midpoint sensor in the housing tube^22^, overestimating actual sleep time^17^. The ethoscope was developed to unequivocally identify immobility periods and assess sleep^23^. We confirmed that social interaction also induced animals to sleep more^15^ (fig 2A). To ensure consistency with previous works^17^, we focused on the first four hours after lights ON, where the effect is unambiguous and reproducible (i.e. 24 h-28 h time window, ZT0-ZT4, fig 2A). Sleep quantification shows a significant difference between social-enriched and lonely animals in this 24 h-28 h period (fig 2B), in line with previous publications^15,17^. Furthermore, *rut* mutant animals showed no significant difference in 24 h-28 h sleep time (fig 2A, B), although socialized *dnc* mutants did exhibit a significant sleep difference in the 24 h-28 h, (fig 2C). Intriguingly, memory-related mutant flies sleep considerably more than their wt counterparts, suggesting additional levels of sleep regulation related to cAMP signaling (fig 2A-C).

**Figure 2.**
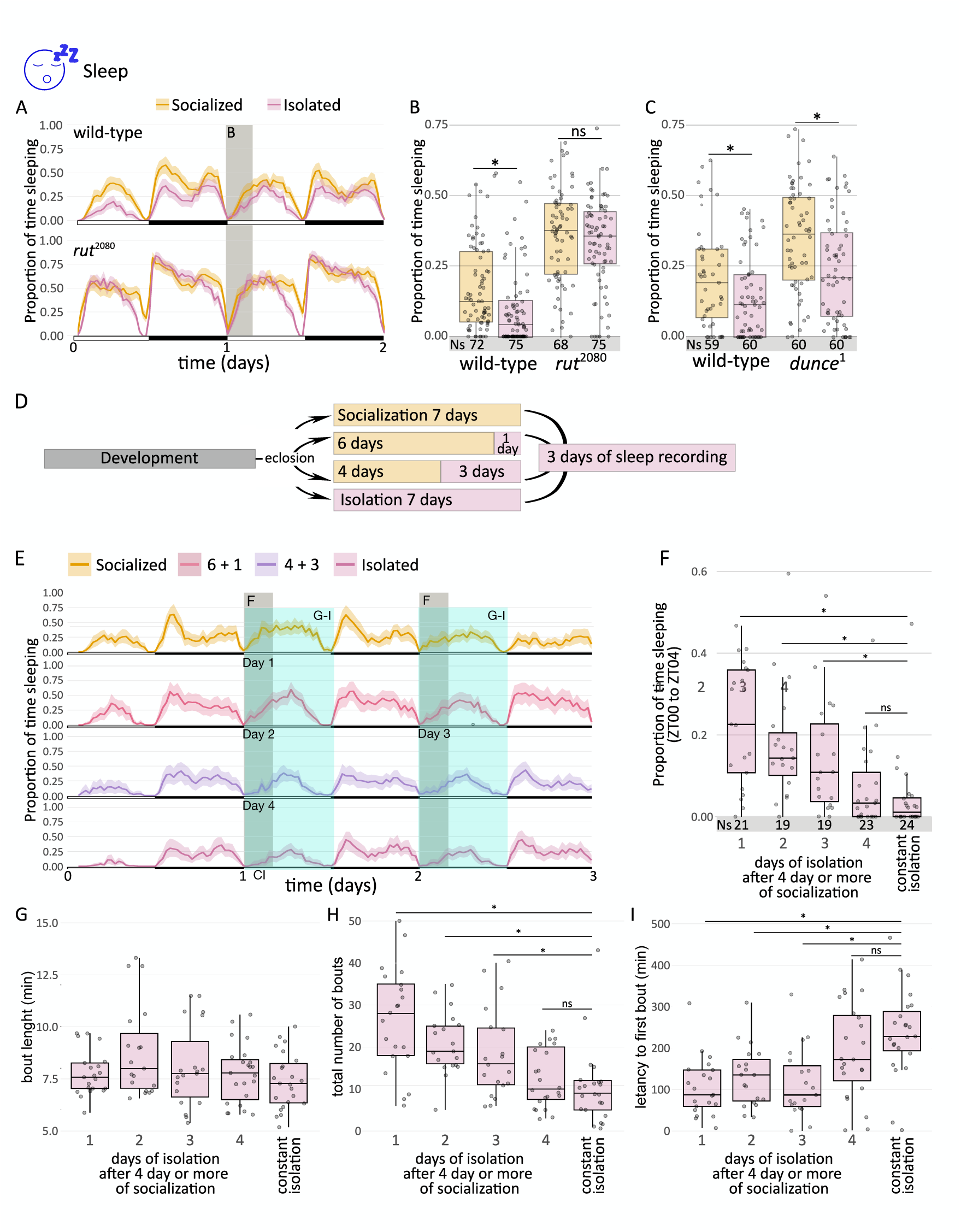
Sleep behavior after socialization is long-term and relies on *rut*. (A) Sleep profile and (B) sleep quantification of the 24 h-28 h time window for *wt* and *rut* mutant background (Kruskal-Wallis chi-squared = 94.165, df = 3, p < 2.2e-16; *post hoc* Dunn comparisons: wt^social^|wt^isolated^ p = 1.07e-02, *rut*^social^|*rut*^isolated^ p = 0.438). (C) Sleep quantification of the 24 h-28 h time window for *wt* and *dnc* mutant background (Kruskal-Wallis chi-squared = 36.476, df = 3, p-value = 5.94e-08; *post hoc* Dunn comparisons: wt^social^|wt^isolated^ p = 3.43e-02, dnc^social^|dnc^isolated^ p = 3.48e-03). (D) Scheme of the sleep time course: flies were either isolated or grouped after eclosion for 7, 6 or 4 days and subsequently isolated for 0, 1 or 3 days (named as socialized, 6+1, 4+3 and constant isolation); after introducing them in ethoscopes, sleep behavior was recorded for 3 days. (E) Sleep profile of animals isolated for 1 to 4 days, using isolated flies as control. Total number of days in isolation for E-I is depicted in the panel. (F) Quantification of sleep from ZT0 to ZT4 for day 1-4 and flies under constant isolation (CI). (G-I) Analysis of bout length (G), total number of bouts (H) and latency to first bout (I) from ZT0 to ZT12 for day 1-4 and animals under CI.

Our results with *dnc* and *rut* mutant flies were apparently contradictory with those previously described using DAMs^15^. However, the ethoscope offers the possibility of analyzing data as they were extracted from DAMs, thus depicting comparable results to published DAM data. This *virtual* DAM analysis did render a significant difference between *rut* mutant grouped and single-reared flies, whereas no sleep changes were apparent in *dnc* mutant animals, in agreement with previous studies (fig S1)^15^. The differing results obtained depending on the type of analysis (regular or virtual DAM) stem from the higher sensitivity of ethoscopes to movement. It also explains why the increased sleep behavior of memory-mutant flies remained unnoticed until now, given that DAMs cannot detect such changes (^24^ and fig S2). Nevertheless, in either case, our data and previous work support the idea that cAMP regulation, necessary for synaptic plasticity, is needed to sustain long-lasting changes in sleep triggered by socialization awareness.

We wondered how long animals that experienced a socially-enriched environment maintained such sleep behavior in absence of social interaction. To compare animals of the same age, we socialized flies for 7, 6 or 4 days (which is enough socialization time in order to generate a sleep effect ^15^) and subsequently isolated them for 0, 1 or 3 additional days (named as socialized, 6+1 or 4+3, respectively). Continuously isolated animals were used as control. Then, their sleep behavior was recorded for the following 3 days (i.e. depicted in fig 2D). In the framework of this experimental approach, we could compare continuously isolated flies with animals isolated for 1 to 4 days after socialization (fig 2E). We could observe a progressive reduction of sleep time in the ZT0-ZT4 after isolation, with significant decrease at day 4 that was comparable to continuous isolation (fig 2F). Thus, 4 days of isolation are enough to modify sleep reaching similarly sleep levels than socially naive flies, in contrast to the need of 5 days described previously using DAMs^17^. The ethoscope also allows a detailed sleep analysis regarding bout length, the total number of bouts and the latency to first bout in a 12-h analysis (ZT0-ZT12). There were no differences in the sleep bout length amongst experimental groups. In contrast, isolated flies for 4 days reduced the number of sleep bouts to similar levels than the ones from socially naive animals, despite we noticed a progressive reduction but still statistically significative (fig 2H). Intriguingly, the latency to the first bout in grouped flies remained similar up to day 3, where it raised sharply, similar to the latency of isolated flies (fig 2I). We conclude that the effect of socialization lasts at least for 3 days, and indeed, it can be considered as long-term.

### SOCIALIZATION-INDUCED DIMINISHED AGGRESSIVENESS DEMANDS cAMP SIGNALING

In isolated flies that previously experienced social interaction, isolation signals starvation and, as a consequence, increases feeding and decreases sleep, meaning that both behavioral changes are reciprocally related^17^. We wondered if socialization awareness was also evident in a different social behavior. Previous data showed that 5-day grouped male flies since eclosion were less aggressive than their single-reared counterparts when tested immediately after the treatment^25^. This behavior also allowed us to determine the progression of long-lasting effects, in order to compare the temporal requirements of social interaction with those of classical learning and memory assays. Thus, we employed animals from both experimental conditions and then evaluated aggression after different isolation periods in a well-established paradigm^26^. Socially-experienced flies showed reduced aggression (i.e. measured as the proportion of time lunging) at 1, 4, 8 and 24 hours after isolation when compared with single-reared animals (fig 3), evidencing a behavioral change at short- and long-term. Despite social interaction had ceased up to 24 h before, grouped flies still spent considerably less time fighting than lonely flies (fig 3), confirming that socialization awareness is a general feature of socialization. Critically, *rut* mutant flies did not show any difference according to their previous experience (fig 3), reinforcing the role of memory-related genes. Besides, our data revealed that at short-term (one hour), grouped *rut* mutant animals were significantly less aggressive than *rut* single-reared flies, thus differentiating from *rut* requirements in classical learning assays. However, this rut-independent effect disappeared after four hours, suggesting that social interaction imprinted a temporary effect independently of cAMP, but long-term socialization consequences in aggression depends on *rut* activity. Overall, *rut* mutant flies showed decreased levels of aggression, which makes comparison with *wt* animals difficult, as previously noticed^27^.

**Figure 3.**
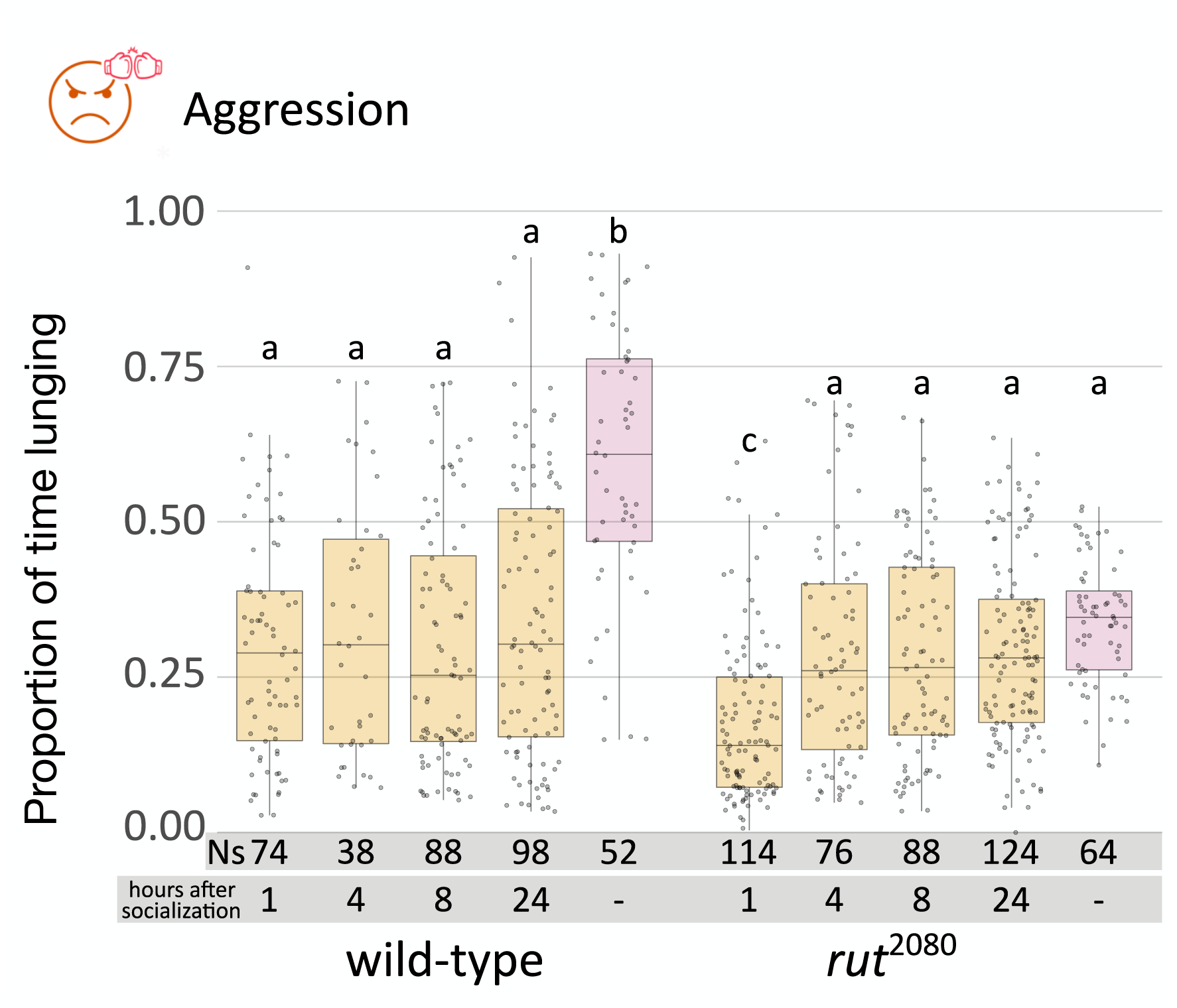
Socialized-reduced aggression shows short- and long-term effect. Quantification of proportion of time expend lunging after different times of re-isolation. Flies either *wt* or in a *rut* mutant background were grouped or isolated for 5 days and then socialized flies were tested after 1, 4, 8 or 24 hours after isolation (Kruskal-Wallis chi-squared = 139.99, df = 9, p-value < 2.2e-16, *post hoc* Dunn comparisons: wt^24h_after_social^|wt^isolated^ p = 1.08e-07, wt^8h_after_social^|wt^isolated^ p = 1.61e-10, wt^4h_after_social^|wt^isolated^ p = 1.88e-05, wt^1h_after_social^|wt^isolated^ p = 2.24e-09, rut^24h_after_social^|rut^isolated^ p = 1.00, rut^8h_after_social^|rut^isolated^ p = 0.815, rut^4h_after_social^|rut^isolated^ p = 0.598, rut^1h_after_social^|rut^isolated^ p = 7.52e-10).

### ANESTHESIA ABOLISHES SOCIALIZATION EFFECTS

Anesthesia blocks long-term memory consolidation in most species^28,29^. In *Drosophila*, a 2-min cold shock acts as anesthetics and is able to impede long-term memory in the classical aversive olfactory conditioning assay^30^. We wondered if anesthesia was also able to block socialization awareness. We exposed adult flies to 3-min cold shock two times per day to single and grouped flies during the training period (fig 4A). Both experimental “cold-shocked” groups did not show any significant differences in food consumption in the 24 h-48 h time window after isolation, in contrast to non-shocked control animals (fig 4B). Given the reciprocal relationship between feeding and sleep behavior regarding social interaction^17^, we confirmed that sleep between lonely and socialized animals in the 24 h-28 h time window also remained similar after cold shock (fig 4C, D). As expected, in non-shocked animals the difference was statistically significant (fig 4C, D). In summary, we found that socialization awareness relies on cAMP signaling and is blocked by anesthesia, as it occurs in long-term memory.

**Fig 4.**
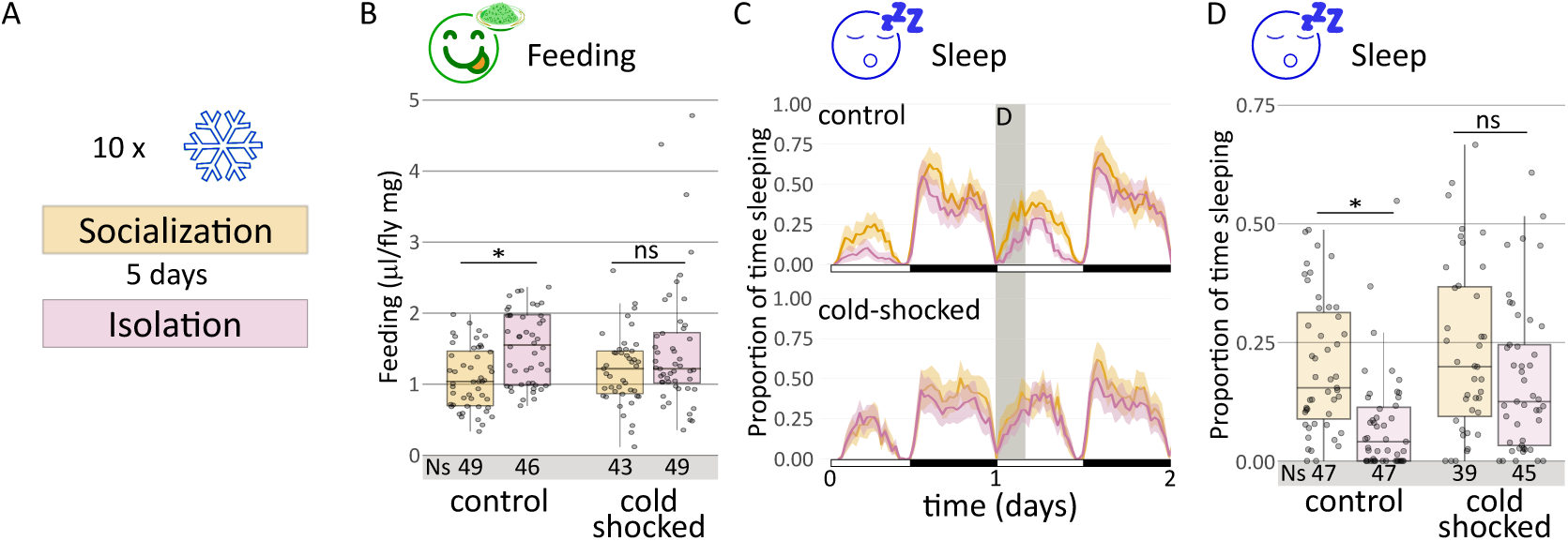
Anesthesia abolishes socialization effects on sleep and food consumption. (A) Scheme of the cold-shock protocol (twice per day). (B) Quantification of food consumption using sCAFE (Kruskal-Wallis chi-squared = 15.954, df = 3, p-value = 1.16e-3; *post hoc* Dunn comparisons: non-shocked^social^|non-shocked^isolated^p = 5.26e-3, shocked^social^|shocked^isolated^ p = 1.00). (C) sleep profile and (D) sleep quantification of the 24 h-28 h time window (Kruskal-Wallis chi-squared = 31.184, df = 3, p-value = 7.78e-07; *post hoc* Dunn comparisons: non-shocked^social^|non-shocked^isolated^ p = 3.05e-06, shocked^social^|shocked^isolated^ p = 0.116) of cold-shocked socialized and isolated *wt* flies, together with non-shocked control *wt* flies.

### SOCIALIZATION CORRELATES WITH INCREASED NEURONAL ACTIVITY AND SYNAPTIC PLASTICITY

In *Drosophila*, LTM increased the number of CREB-activated neurons in the MB^10,31^. To evaluate whether or not socialization also correlates with higher levels of CREB activity in the MB, we used the CAMEL reporter tool after 5 days of socialization directly after eclosion. This tool bears a MB-specific transgenic construct that responds to phosphorylated CREB with the production of GFP^31^. We quantified the number of GFP positive soma (fig 5A) in adult brains, observing an increase in the number of CREB-positive cells in grouped vs single-reared animals (fig 5B). In contrast, this CREB response was lost in *rut* mutant brains (fig 5B).

**Figure 5.**
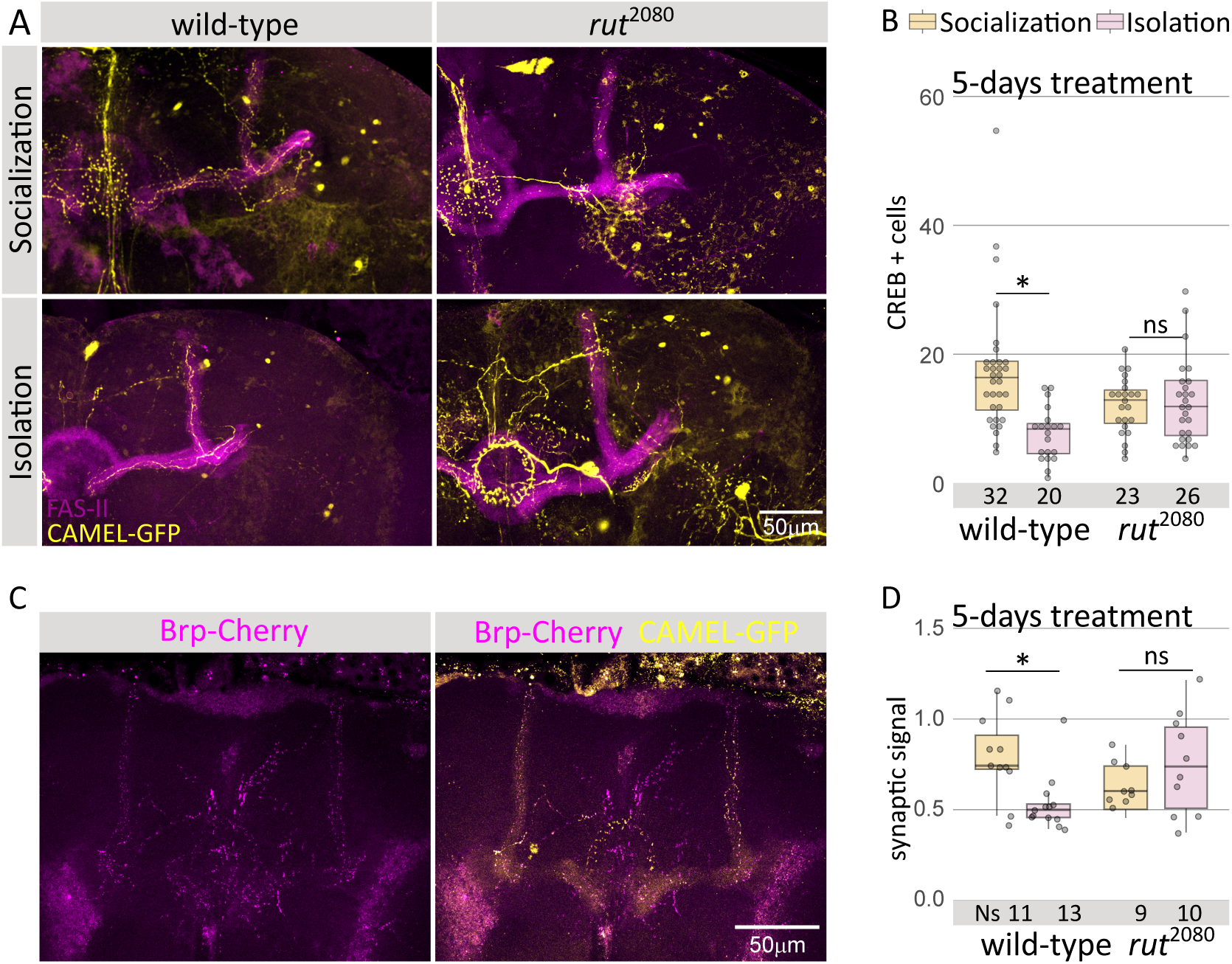
Socialization correlates with cellular and synaptic plasticity. A) Representative confocal images of CAMEL tool for wt and *rut* mutant MB, either socialized or isolated. Only one representative MB is shown. B) Number of CREB GFP-positive cells in the MB of socialized or isolated wt and *rut* mutant animals after 5 days of socialization. Kruskal-Wallis chi-squared = 33.735, df = 3 p-value = 1.93e-05; *post hoc* Dunn key comparisons: 5 days: wt^social^|wt^isolated^ p = 6.01e-05, *rut*^social^|*rut*^isolated^ p = 1.00. C) Example of CAMEL tool (MB cells marked by GFP) combined with the pre-sinaptic marker brp-cherry after 5 days of socialization for wt and *rut* mutant animals, either socialized of isolated. D) Quantification of the number of synapses after either isolation or socialization in both wt and *rut* mutant flies (see fig S5 for a detail on the quantification). Kruskal-Wallis chi-squared = 9.7691, df = 3, p-value = 0.021; *post hoc* Dunn key comparisons: wt^social^|wt^isolated^ p = 2.51 e-02, *rut*^social^|*rut*^isolated^ p = 1.00.

LTM formation using an appetitive conditioning paradigm increased the number of MB-input synapses^32^. Thus, to determine if CREB-activated neurons after socialization also showed signals of increased synaptic plasticity, we included in the CAMEL tool a second reporter, the presynaptic marker BRP, fused with the RFP-variant cherry. This reporter combination allowed the visualization of the presynaptic densities (fig 5C). We quantified the number of synapses per cell volume in brains of 5-day grouped and single-reared animals (fig S2 shows an example of this quantification technique, see M&M). There was a significant increase in the relative number of pre-synapses in the MB of grouped flies compared to single-reared animals (fig 5D), similar to the synaptic plasticity described in mammals after an experience^33^. In contrast, in a *rut* mutant background we could not detect any difference in the number of MB pre-synapses, which was in agreement with the reduced pre-synapse number in *rut* MB-input neurons after appetitive conditioning^32^(fig 5D). Given that intensity of fluorescence varies greatly depending on the region, for analytical purposes we divided the MB in three areas, alpha, beta and the tip of beta. Interestingly, the former two showed only a marginal increase that did not reach statistical significance, however the tip of the MB concentrated most of the increase (fig S3). In summary, results showed a clear correlation of CREB-activated neurons and increased synaptic plasticity with effective social interaction that is abolished in memory impaired mutants, thus supporting a resemblance between socialization awareness and LTM.

## DISCUSSION

Socialization induces several changes in animal behavior and here we show that such changes are long-lasting, as a result of social interaction experience. Not surprisingly, socialization awareness shows similarities with a long-term memory process: involvement of cAMP signaling and processes of neuronal and synaptic plasticity. However, it presents differences with LTM. For instance, the role of *rut* in short-term behavioral changes seems dispensable, at least for aggression (fig 3A, B). It may indicate that short-term effect is independent of *rut*. A striking peculiarity is its temporal dynamics since it would be hard to distinguish putative learning and consolidation stages during socialization, while in long-term memory paradigms both phases are clearly distinguishable (as, for example, in appetitive or aversive olfactory conditioning).

The classic view on sleep regulation indicates that this behavioral state is regulated by the circadian clock and the internal sleep homeostat^34^, but recent work in many species including *Drosophila* show that sleep regulation goes beyond these two processes and includes temperature, starvation, sexual arousal, and social context, among others^35^. Our data suggest that recalling a past social experience may also regulate sleep in flies (fig 1), similar to what happens with psychophysiological insomnia in humans^36^.

In mammals, social isolation has profound effects on behavior and cognition, which is accompanied by detectable alterations in brain structure and function at several levels^37^. For instance, the hippocampus shows reduced dendritic spine density after either postnatal or juvenile social isolation^38,39^*. The* hippocampus is the main structure related to long-term memory, analog to the insect Mushroom Body^40^. In fact, it was previously described that socialization increases the fiber number in the MB, an increase that is impeded by classic learning mutation such as rutabaga^13,14^. In addition, our results revealed that socialization also induces rut-dependent changes in synaptic plasticity of the previously activated MB neurons. The increased synaptic densities in CREB-positive neurons might be explained by the socialization-induced enhanced sleep, given that sleep loss diminishes pre-synaptic densities in cholinergic neurons, including the MB neurons^41,42^. This is unlikely because despite *rut* mutant animals do sleep much more (fig 2B), their MB presynaptic densities do not reach levels of socialized *wt* animals, although they are higher than in isolated flies (fig 5D). Besides, *rut* mutant flies did not reach enough sleep levels as to restore behavioral plasticity^24^, thus suggesting that *rut* increased synaptic activity might be due to the excess of sleep but it is unable to rescue the effect of social interaction (fig 1-3). This reinforces the idea that socialization awareness may induce behavioral plasticity by similar mechanisms to long-term memory.

An apparent contradictory result was that memory-mutant animals did not behave as expected, i.e., as *wt* socialized flies, resembling more to wt isolated flies (fig 1-3). There are several reasons to explain this presumed inconsistency. The most obvious one is that the basal behavior of *rut* and *dnc* mutant flies are different due to the lack of cAMP signaling, as previously described for aggression^27^. Actually, the reduced basal behavioral response of wt isolated animals correlated with their decreased number of CREB-positive cells and pre-sinapses when compared to rut mutant animals, which showed intermediate levels. We might speculate that isolated animals would also generate an “isolation awareness” that enhanced the effect of isolation by diminishing synaptic plasticity due to a cAMP-related mechanisms and, as a consequence, the behavioral response. Why do isolated flies that were previously socialized behave similar to single-reared flies since eclosion^15^? Actually, chronic isolation displays starvation-like phenotypes in *Drosophila*^17^ and starvation disables aversive long-term memory^43^, probably because increased metabolism in the MB and glia are necessary^44,45^. It might well be that socialization awareness was prevented as a consequence of the starvation signaling, and this would explain the similar phenotypes achieved by isolation after socialization and isolation since eclosion, despite mechanistically they should be different. Indeed, one might hypothesize that rescuing such starvation-like phenotype would reveal differences between both experimental conditions.

Notably, socialization-induced behavioral changes are sexually dimorphic, since grouped and single-reared females behave similarly^46^. In fact, male-specific P1 interneurons act as an internal state regulatory hub for sleep, aggression, sleep and spontaneous locomotion^47^. Together with *Diuretic hormone 44*-(*DH44*) and *Tachykinin*-(*TK*) expressing interneurons, P1 neurons form a male-specific neural circuit that regulates spontaneous locomotion in response to social interaction, thus suggesting a possible common mechanism for socially-induced behavioral changes^46^. Interestingly, P1 neurons directly activate a specific subset of dopaminergic neurons that innervate the MB and it drives LTM appetitive olfactory memory formation^48^. The MB is not only a memory regulatory center but also acts as a sleep and feeding regulatory center^49,50^. In this work we have shown that social interaction correlates with increased synaptic plasticity in the MB itself (fig 4). Thus, it is tempting to postulate that socialization awareness may use a general neural circuit connecting P1 neurons, dopaminergic neurons and the MB in order to modify several behaviors with long-lasting effects.

## MATERIALS

### Stocks and fly husbandry

Flies were raised and experiment performed using standard food at 25°C on a 12/12h light/dark cycle. *rutabaga^2080^* (#9405), *dunce*^*1*^ (#6020) and *Wild Type* (*Canton S* #64349) stocks were obtained from Bloomington Drosophila Stock Center. The CAMEL tool is composed by 6xCRE-splitGal4^AD^, UAS-eGFP and R21B06-splitGal4^DBD^, gently donated by Dr Jan Pielage^31^. *rut^2080^; 6xCRE-splitGal4^AD^* and *UAS-cherry-Brutchpilot; R21B06-splitGal4^DBD^* stocks were combined in our laboratory and are available under request.

### Isolation/socialization Protocol

Male virgin flies were collected under CO_2_ anesthesia within 4 hours post-eclosion and isolated in individual glass vials or socialized (25:25 male:female) in a plastic bottle. After 5 days of socialization or isolation, all flies were isolated without using anesthesia for 24 h (except where indicated) and then, behavioral experiments or dissections were performed.

In the case of cold shock, flies were ice-cold shocked twice a day (Zeitgeber Time 1 -ZT01- and ZT9) during the five days of isolation/socialization protocol for 2-3 minutes (i.e. until flies fainted). Glass vials were used to allow good cold transfer from ice. Afterwards vials were placed horizontally in a RT surface to let flies recover.

### single fly Capillary Feeding (sCAFE)

The protocol from ^18^ was used with slight modifications. Males were placed in individual vials with a wet filter paper at the bottom and a 5 μl capillary (Blaubrand, 708707) with 5% sucrose water food. The capillary was introduced through a 5mm cut 200 μl pipet tip that goes through a wet plug and sustained with an additional tip. After 24 h food intake is measured (0 h-24 h time window), the capillary substituted by a new one and plugs are wet again to preserve moisture. 24 h later food intake is measured again. Once the experiment has finished flies are weighted. Additional 3 individual tubes without flies were measured to control the evaporation rate.

### Sleep

For all experiments, flies were sorted into glass tubes [70 mm × 5 mm × 3 mm (length × external diameter × internal diameter)] containing the same food used for rearing under a regime of 12:12 Light:Dark (LD) condition in incubators set at 25°C. Activity recordings were performed using ethoscopes^23^. Behavioural data analysis was performed in RStudio (RStudio Team. RStudio: Integrated Development for r. RSudio, Inc. Boston, MA; 2015. http://www.rstudio.com/) employing the Rethomics suite of packages^51^. All sleep assays were repeated at least twice with 20–40 flies/treatment/experiment.

### Aggression

The protocol from^26^ with slight modifications was used. Briefly, two flies were placed into each chamber of the arena (4×3 mm grid) with food. One-to-one socialization was achieved by allowing both flies to interact, whereas isolation was caused by a black divider that allowed physical separation of flies. After 5 days, socialized flies were also separated by the divider for 1, 4, 8 or 24 h. After removing the divider, reunited flies were recorded for 20 minutes and agression analyzed by means of the *FlyTracker* (MATLAB) software and the platform JAABA (*Janelia Automatic Animal Behavior Annotator*), that identifies when the animal is lunging. The proportion of time fighting is the number of frames in which a particular animal lunges divided by the total number of frames.

### Immunolabeling, imaging and image analysis

Adult brain preparations were stained following the same protocol as in^52^. Dissections were always performed at ZT4-5 to avoid possible circadian-induced changes.

For CREB+ cells experiment, primary antibodies used were anti-GFP rabbit (1/200; DSHB???REF) and anti-Fasciclin II mouse (1/50; DSHB). To quantify synapse number, primary antibodies used were anti-GFP goat (1/200; DSHB), anti-RFP rabbit (1/200; DSHB) and anti-Fasciclin II mouse (1/50; DSHB). Secondary antibodies used were Alexia 488, 568 or 680 (1/500; Life Technologies). Images were taken by a Leica SP5 confocal microscopy re-using the same experimental conditions, avoiding saturation. CREB+ cell images were taken using a 40X objective, with slices of 3 um. Synapse quantification confocal images were taken the same day using a 63X objective, slices of 0,8um. Posteriorly images were treated using Imaris 6.3.1 software. Axon volume was rebuilt using the Volume tool and brutchpilot signal was quantified using the Spots tool. To adjust brightness parameters accurately the MB was divided in three parts (alfa, beta and beta tip) (Fig. Supp. 2). Synaptic density for each Mushroom Body is the summatory of spots/volume from each part.

### Statistical analysis

For the behavioral and morphological experiments (figures 1, 2, 3, 4, 5, S1 and S3), the data was analyzed in R (version 3.6.3) through Rstudio (Version 1.0.153), employing the Kruskal-Wallis non-parametric test (library *stats*). When appropriate, we performed post hoc Dunn analyses (library FSA) to identify specific differences between treatments. All assays were repeated at least twice with sample sizes as indicated within the figure

## ACKNOWLEDGEMENTS

We would like to thank Javier Gil Castillo for his invaluable help and advices in 3D printing. We also appreciate flies and reagents from the Bloomington and VDRC stock centers. Special thanks to our colleagues Prof Alberto Ferrús, Dr Sergio Casas-Tintó, Dr Pablo Méndez, Dr Abhijit Das and Dr JL Trejo-Pérez for their helpful comments and suggestions on this manuscript. Special thanks to Dr Pavan Agrawal and his lab for their critical reading and suggestions on BioRxiv manuscript. We thank the support of the scientific image and microscopy unit (Cajal Institute).

FAM was a recipient of a RyC-2014-14961 contract (2016-2022). Grant RyC-2014-14961 funded by MICIU/AEI/10.13039/501100011033 and by ESF Investing in your future. Grant CNS2022-135223 funded by MICIU/AEI/10.13039/501100011033 and by European Union NextGeneration EU/PRTR.

BG-M is a recipient of a FPI-UAM predoctoral fellowship, grant number SFPI/2020/00878. JIM was a recipient of a JAE intro fellowship (grant number JAEINT_22_01271) funded by the Spanish National Research Council (CSIC).

EJB is a member of the Argentine Research Council (CONICET), and he is funded by *Agencia Nacional de Promoción de la Investigación, el Desarrollo Tecnológico y la Innovación*, Argentina, through grants PICT-2020-SERIEA-01240, PICT-PRH-2021-00009, and CONICET through grant PIP 11220200102510CO.

## Author contributions

FAM conceptualized and designed the project; FAM and EJB supervised the project; BGM, GT and JIM performed aggression experiments and cellular studies; BGM and APZ performed feeding assays; GSAT and EJB performed and analyzed sleep experiments; EJB and ET analyzed data; FAM and EJB wrote the original draft and made the figures; ET made extensive editing to the manuscript and revised statistics; FAM and EJB were responsible of funding acquisition.

## Competing interests

The authors declare no competing interests.

## Notes

### Competing Interest Statement

The authors have declared no competing interest.

### Summary of Updates

Authorship updated; text (results and discussion) and fig 3 revised.

