## Supplementary material for "Socialization causes long-lasting behavioral changes": Fig S1-S3

**SUPPLEMENTARY INFORMATION**

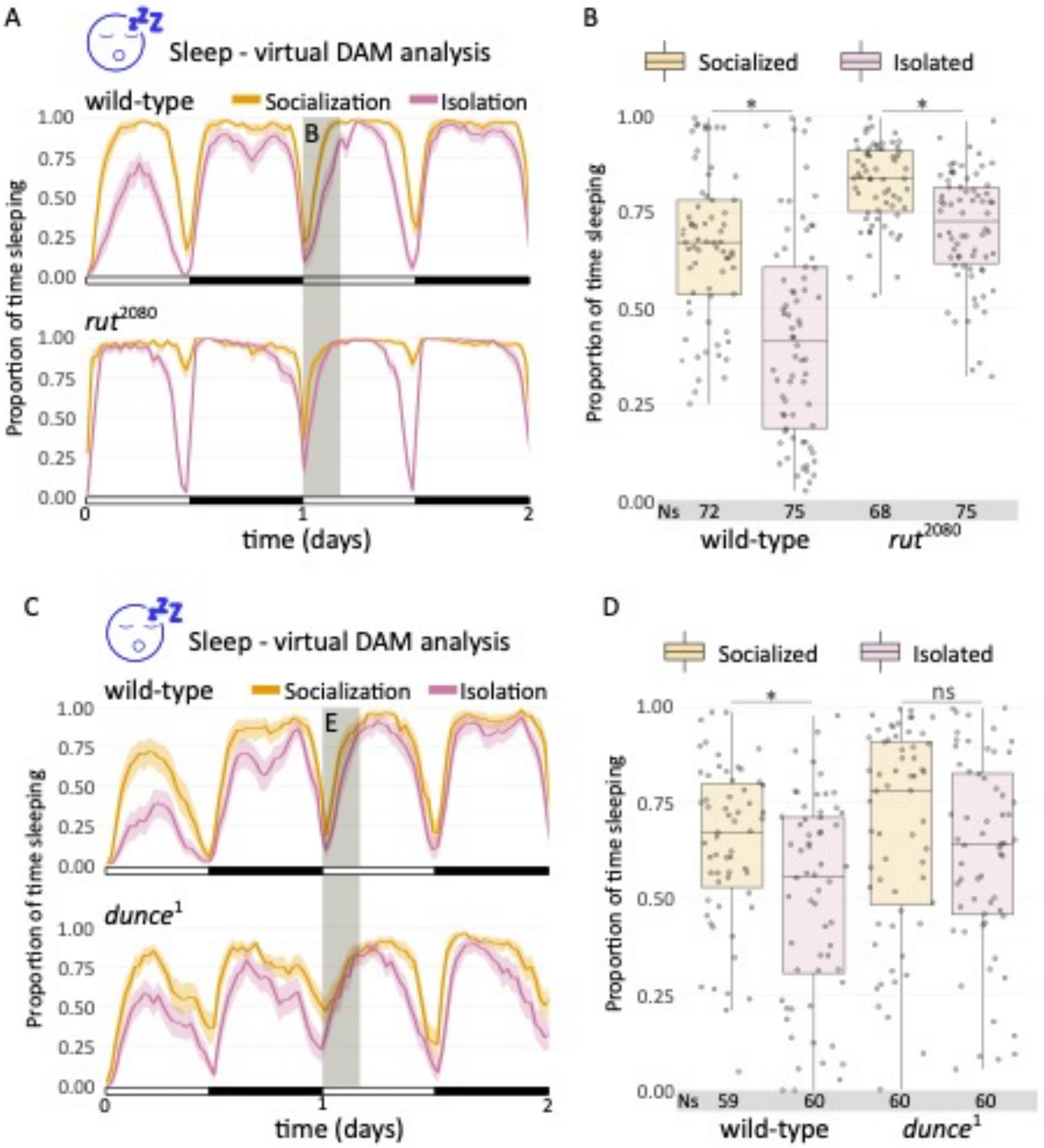


**Figure S1. Ethoscope and Virtual DAM sleep analyses show significant differences**. A) Virtual DAM sleep profile and (B) virtual DAM sleep quantification of the 24-28 h window of *wt* and *rut* mutant flies in socialized and isolated conditions obtained with ethoscopes (Kruskal-Wallis chi-squared = 86.153, df = 3, p-value < 2.2e-16; *post hoc* Dunn comparisons: $wt^{social}|wt^{isolated}$ p = 1.03e-05, $rut^{social}|rut^{isolated}$ p = 3.13e-04). C-D) Analysis of *wt* and *dnc* mutant flies in socialized and isolated conditions. C) Virtual DAM sleep profile and (D) virtual DAM sleep quantification of the 24-

28 h window (Kruskal-Wallis chi-squared = 15.606, df = 3, p-value = 0.001; *post hoc* Dunn comparisons: $wt^{social}|wt^{isolated}$ p = 0.025, $dnc^{social}|dnc^{isolated}$ p = 0.160).

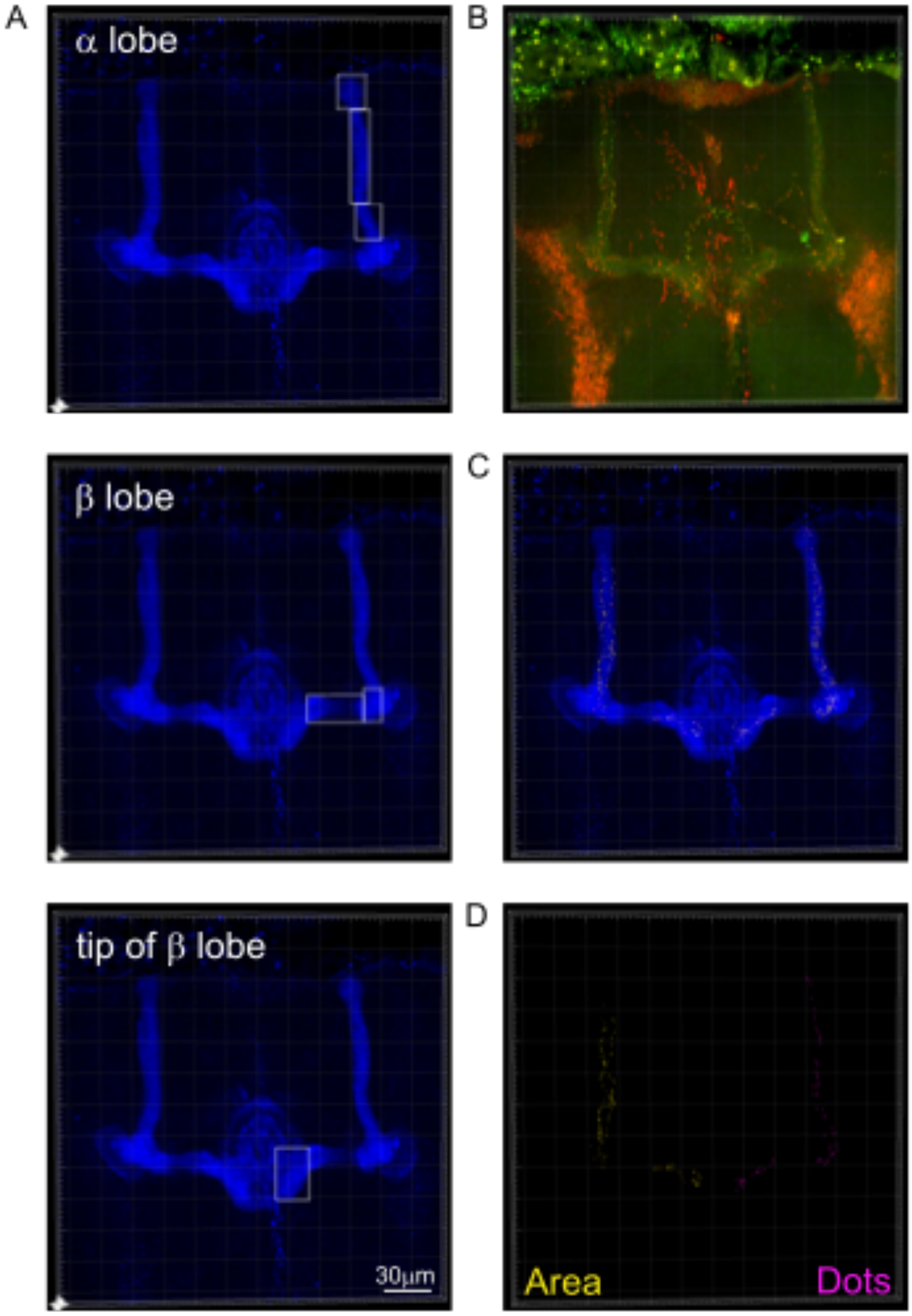


**Figure S2. IMARIS treatment of confocal images obtained by CAMEL tool combined with bruchpilot cherry.** A) The first step is to segment the region of interest. Given the differences in brightness, we decided to segment each MB into 3 regions representing alfa (3), beta (2) and tip of beta (1) lobes . B) Original .TIFF image from the confocal microscope depicting GFP from CAMEL in green and BRP in red. C) After selecting the region of interest, we selected the fluorescence channels (GFP for volume, cherry for spots-active zones). In the case of Volume, we used "Absolute intensity thresholding" unchecking Smooth in order to identify the cell volume, removing small particles (number of voxels filter) and getting the Volume Sum. For Spots, we estimated the XY diameter of the synapsis (in our case, 0.42), unchecking background subtraction, using Intensity Mean from the Volume Channel (to render Spots exclusively within the neuronal Volume) and getting the Total Number of Spots. We divided the number of Spots by the Volume. D) Reconstruction of Volume (left, yellow) and Spots (right, purple) created by IMARIS software.

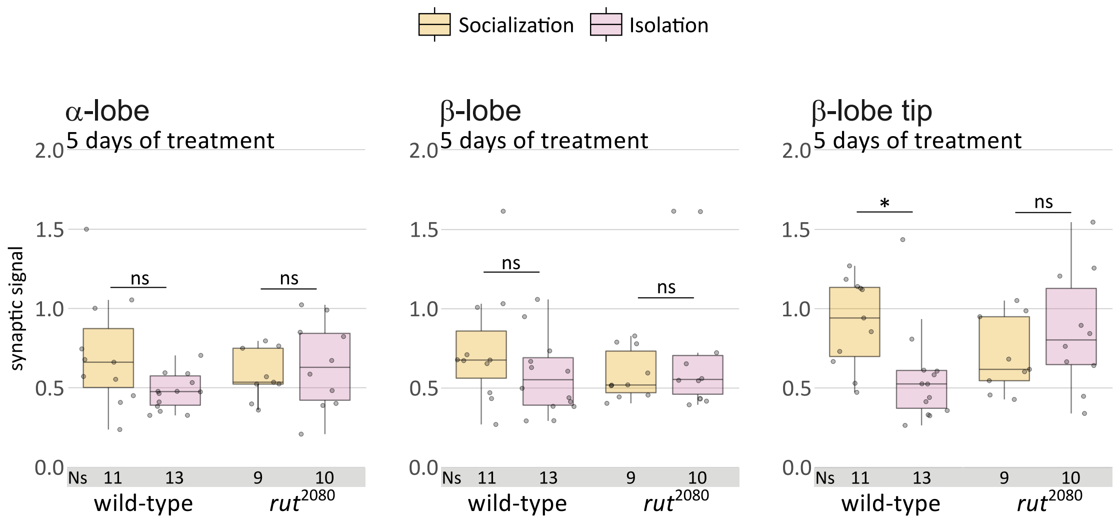


**Figure S3. Regional quantification of the synapsis number in the MB.** α, β and β tip lobe quantification for wt and *rut* mutant flies, as shown by the scheme of fig S2. α-lobe: Kruskal-Wallis chi-squared = 3.7824, df = 3, p-value = 0.286; β-lobe: Kruskal-Wallis chi-squared = 2.2608, df = 3, p-value = 0.520; β-lobe tip: Kruskal-Wallis chi-squared = 9.7365, df = 3, p-value = 0.021, *post hoc* Dunn comparisons: $wt^{social}|wt^{isolated}$ p = 0.022, $rut^{social}|rut^{isolated}$ p = 0.149.
